# Physiological relevance, localization and substrate specificity of the alternative (type II) mitochondrial NADH dehydrogenases of *Ogataea parapolymorpha*

**DOI:** 10.1101/2021.04.28.441406

**Authors:** Hannes Juergens, Álvaro Mielgo-Gómez, Albert Godoy-Hernández, Jolanda ter Horst, Janine M. Nijenhuis, Duncan G. G. McMillan, Robert Mans

**Affiliations:** Department of Biotechnology, Delft University of Technology, Delft, The Netherlands

**Author notes:** Corresponding authors: Robert Mans, and Duncan McMillan.

## Abstract

Mitochondria from *Ogataea parapolymorpha* harbor a branched electron-transport chain containing a proton-pumping Complex I NADH dehydrogenase and three alternative (type II) NADH dehydrogenases (NDH2s). To investigate the physiological role, localization and substrate specificity of these enzymes, growth of various NADH dehydrogenase mutants was quantitatively characterized in shake-flask and chemostat cultures, followed by oxygen-uptake experiments with isolated mitochondria. Furthermore, NAD(P)H:quinone oxidoreduction of the three NDH2s were individually assessed. Our findings show that the *O. parapolymorpha* respiratory chain contains an internal NADH-accepting NDH2 (Ndh2-1/OpNdi1), at least one external NAD(P)H-accepting enzyme and likely additional mechanisms for respiration-linked oxidation of cytosolic NADH. Metabolic regulation appears to prevent competition between OpNdi1 and Complex I for mitochondrial NADH. With the exception of OpNdi1, the respiratory chain of *O. parapolymorpha* exhibits metabolic redundancy and tolerates deletion of multiple NADH-dehydrogenase genes without compromising fully respiratory metabolism.

**Importance:** To achieve high productivity and yields in microbial bioprocesses, efficient use of the energy substrate is essential. Organisms with branched respiratory chains can respire *via* the energy-efficient proton-pumping Complex I, or make use of alternative NADH dehydrogenases (NDH2s). The yeast *Ogataea parapolymorpha* contains three uncharacterized, putative NDH2s which were investigated in this work. We show that *O. parapolymorpha* contains at least one ‘internal’ NDH2, which provides an alternative to Complex I for mitochondrial NADH oxidation, albeit at a lower efficiency. The use of this NDH2 appeared to be limited to carbon excess conditions and the *O. parapolymorpha* respiratory chain tolerated multiple deletions without compromising respiratory metabolism, highlighting opportunities for metabolic (redox) engineering. By providing a more comprehensive understanding of the physiological role of NDH2s, including insights into their metabolic capacity, orientation and substrate specificity this study also extends our fundamental understanding of respiration in organisms with branched respiratory chains.

## Introduction

Dissimilation of glucose to carbon dioxide results in the generation of reducing equivalents in the form of NADH, which are continuously (re)oxidized and intrinsically linked to the formation of ATP by the respiratory chain. In eukaryotes, glucose dissimilation and NADH generation occurs in the cytosol by glycolysis and in the mitochondrial matrix by the combined action of the pyruvate-dehydrogenase complex and the tricarboxylic acid cycle. NADH cannot cross the inner mitochondrial membrane (1), and as a result needs to be (re)oxidized in the cellular compartment in which it is generated.

Fungal respiratory chains are branched and contain multiple entry points for electrons from NADH. While the type I NADH:quinone oxidoreductase (NDH1 or Complex I) couples oxidation of mitochondrial NADH to the translocation of protons over the inner mitochondrial membrane, most fungi also possess so-called alternative NADH dehydrogenases (NDH2s) that catalyze NADH:quinone oxidoreduction without proton translocation (2). These ‘alternative NADH dehydrogenases’ are monotopic proteins that attach to the inner mitochondrial membrane, but may differ in which side of the membrane they are located. Their catalytic sites either face the mitochondrial matrix (‘internal’) where their catalytic activity overlaps with Complex I, or the intermembrane space (‘external’), allowing direct oxidation of cytosolic NADH (3–5). A prominent example for the phenotypical role of NDH2 can be found in the yeast *Saccharomyces cerevisiae*, which does not harbor a Complex I and instead relies solely on one internal and two external alternative NADH dehydrogenases as entry points for NADH-derived electrons into the respiratory chain (6, 7). Besides NADH, some external fungal NDH2s, such as the external alternative dehydrogenases from *Kluyveromyces lactis* (8, 9) and *Neurospora crassa* (10–12), have also been reported to accept NADPH as substrate, either exclusively, or in addition to NADH. Based on the distribution of a key acidic residue (E272 in Ndi1 from *S. cerevisiae*), which has been proposed to prevent interaction with the phosphate group of NADPH (13, 14), it has been suggested that the majority of NDH2s oxidize NADH rather than NADPH (15).

*Ogataea parapolymorpha* (formerly *Hansenula polymorpha*) is a methylotrophic, thermotolerant, Crabtree-negative yeast that is characterized by its rapid aerobic growth (16, 17). It has a branched respiratory chain that contains Complex I and three putative alternative NAD(P)H dehydrogenases (referred to as Ndh2-1, Ndh2-2 and Ndh2-3 in this study) with unknown substrate specificity and unknown orientation on the inner mitochondrial membrane (18). Interestingly in the presence of excess glucose and oxygen, deletion of Complex I from *O. parapolymorpha* does not result in a reduction of the maximum specific growth rate or biomass yield. This implies that Complex I is disposable under these conditions. In aerobic glucose-limited cultures, elimination of Complex I resulted in mutant strain which exhibited a 16% lower biomass yield while it maintained a fully respiratory metabolism (18). These phenotypes suggested that *O. parapolymorpha* harbors at least one internal alternative NADH dehydrogenase capable of compensating for the lack of Complex I, albeit at a lower efficiency of respiration-coupled ATP production. Such an internal enzyme also would likely be responsible for (re)oxidation of mitochondrial NADH under fast-growing glucose-excess conditions, especially since NDH2s appear to be suited to catalyze the high rates of NAD^+^ regeneration required to sustain a high glycolytic flux in the absence of fermentation (19).

The aim of this study was to investigate the physiological roles of the three putative alternative NAD(P)H dehydrogenases of *O. parapolymorpha* with a special focus on determining whether this yeast indeed possesses an internal alternative NADH dehydrogenase capable of functionally substituting Complex I. To this end, the aerobic growth characteristics of various *O. parapolymorpha* NAD(P)H-dehydrogenase mutant strains were investigated in glucose-grown batch- and chemostat cultures. To determine localization and substrate specificity of the alternative dehydrogenases, substrate-dependent oxygen-uptake rates of isolated mitochondria from wild-type and mutant strains were measured, and activity of the individual membrane-bound dehydrogenases with NADH and NADPH was assessed.

## Results

### Disruption of Ndh2-1 leads to a reduction of specific growth rate and a Crabtree-positive phenotype in *O. parapolymorpha*

To investigate the contribution to respiration of the three putative alternative NAD(P)H dehydrogenases, strains IMD003, IMD004 and IMD005 were constructed, which harbored a disrupted version of the structural gene for Ndh2-1, Ndh2-2 or Ndh2-3, respectively. The physiology of these mutant strains was then assessed in aerobic shake-flask batch cultures in the presence of excess glucose (2 g L^−1^). The high specific rate of oxygen uptake by Crabtree-negative yeasts can lead to oxygen limitation in shake-flask cultures resulting in respiro-fermentative metabolism (20). However, under the cultivation conditions in this study the wild-type *O. parapolymorpha* strain CBS11895 grows with a fully respiratory phenotype and at a comparable specific growth rate as previously described in fully aerobic bioreactors, indicating that oxygen limitation did not occur (Table 1) (17).

**Table 1:** Physiology of wild-type *O. parapolymorpha* CBS11895 and congenic mutant strains in aerobic shake-flask cultures grown at 30°C in synthetic medium with urea as nitrogen- and glucose (2 g L^−1^) as carbon source. Data are presented as mean ± mean absolute deviation from at least two independent shake-flask cultures. Specific growth rates and ethanol yield were calculated from the exponential growth phase.

| Strain (genotype) | Specific growth rate (h<sup>-1</sup>) | Physiology |
| --- | --- | --- |
| CBS11895 (*wild type*) | 0.36 ± 0.01 | Respiratory |
| IMD003 ($\Delta ndh2-1$) | 0.30 ± 0.01 | 0.31 ± 0.02 mol<sub>EtOH</sub><br>mol<sub>Glucose</sub><sup>-1</sup> |
| IMD004 ($\Delta ndh2-2$) | 0.34 ± 0.01 | Respiratory |
| IMD005 ($\Delta ndh2-3$) | 0.35 ± 0.01 | Respiratory |
| IMX1978 ($\Delta ndh2-2 \Delta ndh2-3$) | 0.32 ± 0.00 | Respiratory |
| IMX2017 ($\Delta ndh2-1 \Delta ndh2-2 \Delta ndh2-3$) | 0.30 ± 0.00 | 0.29 ± 0.02 mol<sub>EtOH</sub><br>mol<sub>Glucose</sub><sup>-1</sup> |
| IMX2197 ($\Delta nubm \Delta ndh2-2 \Delta ndh2-3$) | 0.31 ± 0.00 | Respiratory |

In shake-flask cultures, the specific growth rate of mutant strain IMD005 *(Δndh2-3)* did not differ significantly from that of the wild-type strain CBS11895, whereas strains IMD003 *(Δndh2-1)* and IMD004 *(Δndh2-2)* exhibited 17 and 6% lower specific growth rates, respectively (Table 1). In cultures of strains CBS11895, IMD004 and IMD005 no fermentation products were detected, indicating that a single disruption of *NDH2-2* or *NDH2-3* did not impede aerobic respiratory metabolism. In contrast, strain IMD003 exhibited a Crabtree-positive phenotype, producing 0.31 ± 0.02 mol of ethanol for each mol of glucose consumed. We then combined disruptions in *NDH2-2* and *NDH2-3*, resulting in strain IMX1978, which exhibited a slightly lower specific growth rate than the strains with individually disrupted *NDH* genes, but still maintained a respiratory metabolism (Table 1). An additional disruption in strain IMX1978 of *NUBM*, which encodes an essential 51 kDA subunit of respiratory Complex I (21), also did not result in detectable ethanol production in the resulting strain IMX2197 *(Δnubm Δndh2-2 Δndh2-3)*. Collectively, these results demonstrate that Ndh2-1 presence alone suffices for supporting NAD(P)H turnover requirements in *O. parapolymorpha* for fully respiratory growth in glucose-grown batch cultures.

Strain IMX2017, in which all three *NDH2* genes were disrupted, exhibited a phenotype similar to that of strain IMD003 *(Δndh2-1)*, displaying a 17% lower specific growth rate than observed for the wild-type strain CBS11895, and an ethanol yield similar to IMD003 (Table 1). Since yeast respiratory chains typically possess either none, or a single internal alternative NAD(P)H dehydrogenase (3), and respiratory Complex I is not physiologically relevant under these conditions in *O. parapolymorpha* (18), these data support the hypothesis that Ndh2-1 has an internal orientation (i.e. facing the mitochondrial matrix).

### Oxygen consumption studies with mitochondria from wild-type *O. parapolymorpha* and deletion mutants confirm internal orientation of Ndh2-1

Measurement of the oxygen-uptake rates of isolated mitochondria using a compartmentalized substrate approach has been previously used to determine the orientation of yeast mitochondrial NAD(P)H dehydrogenases (7, 8, 22). To obtain *O. parapolymorpha* mitochondria, we adapted a protocol for isolation of mitochondria from glucose-limited *S. cerevisiae* cultures (7). Initial isolations from wild-type cells grown in glucose-limited chemostat cultures resulted in well-coupled *O. parapolymorpha* mitochondria with a ‘respiratory control ratio’ (RCR) of 3.6 when assayed with endogenous NADH (generated in the mitochondrial matrix by addition of pyruvate and malate). However, the same preparations exhibited rapid, uncoupled oxygen uptake when exposed to methanol or ethanol. Methanol- and ethanol-dependent oxygen-uptake rates were approximately 4-fold and 2.5-fold faster, respectively, than ADP-stimulated respiration of endogenous NADH. (M)ethanol-dependent oxygen uptake indicated a potential contamination of the mitochondrial preparations of these glucose-derepressed cultures with peroxisomes containing methanol oxidase (MOX) (23, 24). A similar contamination was previously observed in mitochondrial preparations from the methylotrophic yeast *Pichia pastoris* (22). In an attempt to obtain mitochondrial preparations devoid of MOX activity, mitochondria were also isolated from cells grown under MOX-repressing conditions in glucose-grown batch- or nitrogen-limited chemostat cultures (25, 26). Mitochondrial isolations from cells grown under these conditions did not consume oxygen in the presence of methanol, but they exhibited RCRs close to 1 when assayed with endogenous NADH, indicating uncoupled preparations (data not shown). In light of these findings, mitochondria isolated from glucose-limited chemostat cultures were used for respiration studies, and interference of the oxygen-uptake measurements by MOX activity was minimized by using reaction mixtures and substrates devoid of alcoholic solvents (see Methods section).

To test the hypothesis that the catalytic site of Ndh2-1 is oriented towards the mitochondrial matrix, oxygen uptake was measured in the presence of endogenous and exogenous NADH using mitochondria isolated from wild-type *O. parapolymorpha* (CBS11895), from strains possessing only Complex I (IMX2017; *NUBM Δndh2-1 Δndh2-2 Δndh2-3*) or only Ndh2-1 (IMX2197; *Δnubm NDH2-1 Δndh2-2 Δndh2-3*) as known respiration-linked NAD(P)H dehydrogenases (Table 2). These strains were of specific interest because in aerobic glucose-limited chemostats at a dilution rate of 0.1 h^−1^, all three strains exhibited a fully respiratory metabolism (see below, Table 3), indicating a functional respiratory chain responsible for (re)oxidation of both cytosolic and mitochondrial NADH.

**Table 2:**
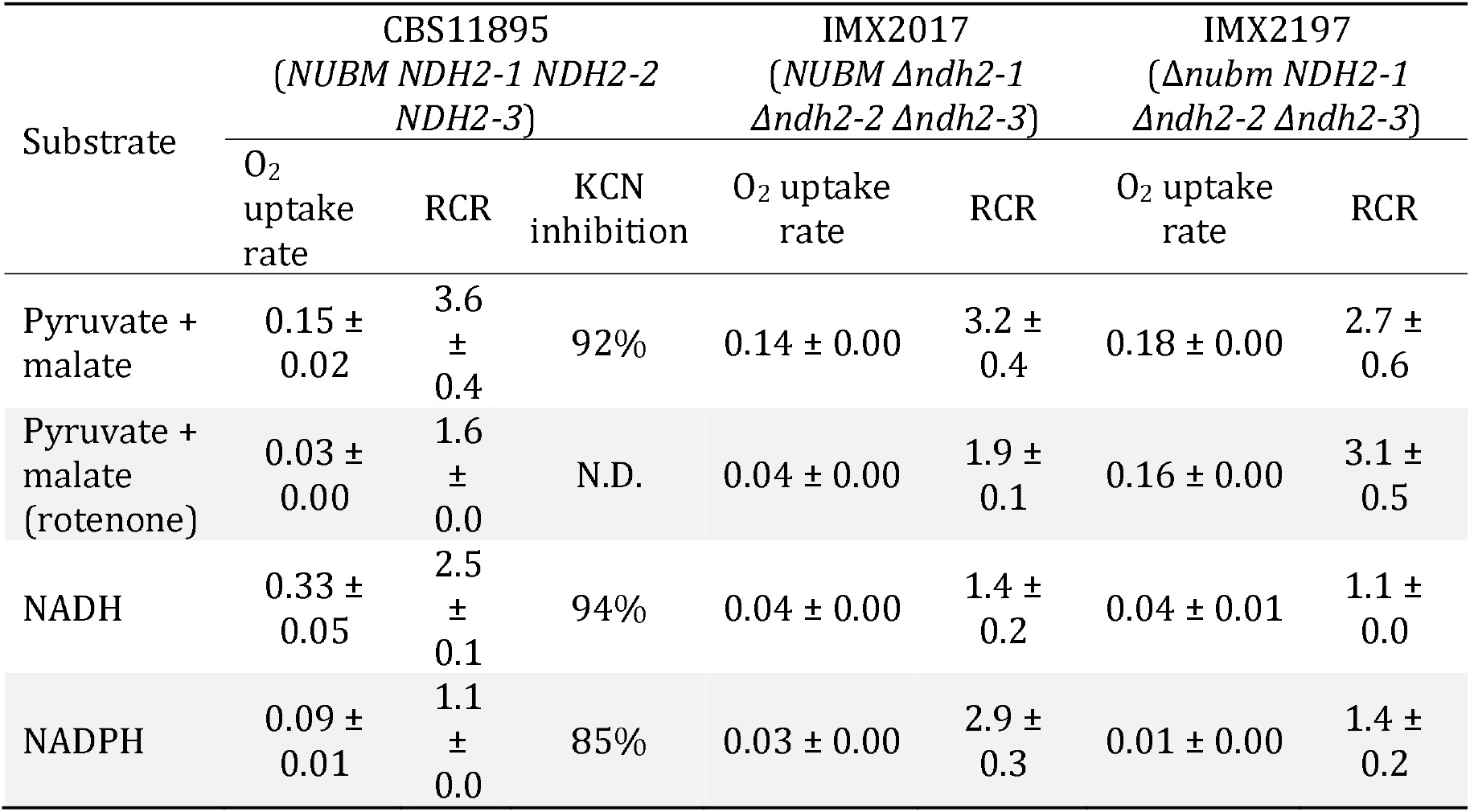
Substrate-dependent rates of oxygen uptake by mitochondria from wild-type *O. parapolymorpha* (CBS11895) and mutants possessing only Complex I (IMX2017) or Ndh2-1 (IMX2197) as known respiration-linked NAD(P)H dehydrogenases. Mitochondria were isolated from cells grown in aerobic, glucose-limited chemostat cultures at a dilution rate (D) of 0.1 h^−1^ and assayed at 30°C, pH 7.0. Oxygen uptake in μmol O_2_ (mg protein)^−1^ min^−1^ was determined in the presence of 0.25 mM ADP. Respiratory control ratio (RCR) values represent the ratio of oxygen uptake rates in the presence and absence (prior to addition) of ADP. For tests with rotenone (50 μM), mitochondria were pre-incubated with this inhibitor at assay conditions prior to substrate addition. Oxygen-uptake rates and RCRs are presented as mean ± standard deviation of measurements with mitochondria from at least two independent chemostat cultures for each strain. Tests with KCN (1 mM) were performed with mitochondria from a single, representative isolation of strain CBS11895. N.D., not determined.

**Table 3:** Physiology of *Ogataea parapolymorpha* strains CBS11895, IMX2017 and IMX2197 in aerobic, glucose-limited chemostat cultures grown at a dilution of 0.1 h^−1^, 30°C and pH 5. Data are presented as mean ± mean absolute deviation from two independent replicates. Carbon recoveries were calculated based on a biomass carbon content of 48% (w/w). Symbols: Y_X/S_ and Y_X/O2_ = yield of biomass dry weight on glucose and oxygen, respectively; RQ = respiratory quotient; q_Glucose_, q_CO2_ and q_O2_ represent biomass-specific uptake/production rates of glucose, CO_2_ and O_2_, respectively; C_X_ represents biomass dry weight concentration. *Data for CBS11895 were reproduced from Juergens, Niemeijer (17).

| Strain | CBS11895*<br>wild type | IMX2017<br>only Complex I | IMX2197<br>only Ndh2-1 |
| --- | --- | --- | --- |
| Actual dilution rate (h<sup>-1</sup>) | 0.10 ± 0.00 | 0.10 ± 0.00 | 0.10 ± 0.00 |
| Reservoir glucose (g L<sup>-1</sup>) | 7.49 ± 0.00 | 7.36 ± 0.01 | 9.04 ± 0.01 |
| Y<sub>X/S</sub> (g biomass [g glucose]<sup>-1</sup>) | 0.51 ± 0.00 | 0.52 ± 0.00 | 0.44 ± 0.01 |
| Y<sub>X/O2</sub> (g biomass [g O<sub>2</sub>]<sup>-1</sup>) | 1.35 ± 0.05 | 1.36 ± 0.00 | 0.94 ± 0.01 |
| RQ (-) | 1.05 ± 0.01 | 1.05 ± 0.00 | 1.04 ± 0.00 |
| q<sub>Glucose</sub> (mmol [g biomass]<sup>-1</sup> h<sup>-1</sup>) | -1.08 ± 0.03 | -1.07 ± 0.01 | -1.28 ± 0.00 |
| q<sub>CO2</sub> (mmol [g biomass]<sup>-1</sup> h<sup>-1</sup>) | 2.44 ± 0.07 | 2.40 ± 0.01 | 3.50 ± 0.00 |
| q<sub>O2</sub> (mmol [g biomass]<sup>-1</sup> h<sup>-1</sup>) | -2.32 ± 0.08 | -2.30 ± 0.01 | -3.38 ± 0.00 |
| C<sub>X</sub> (g biomass L<sup>-1</sup>) | 3.84 ± 0.08 | 3.83 ± 0.04 | 4.01 ± 0.05 |
| Carbon recovery (%) | 99.3 ± 1.2 | 100.1 ± 0.8 | 98.8 ± 0.8 |

Mitochondria isolated from strain CBS11895 readily consumed oxygen in the presence of endogenous and exogenous NADH (Table 2). RCRs for endogenous NADH (3.6 ± 0.4) and exogenous NADH (2.5 ± 0.1) indicated that the observed respiration-linked NADH oxidation is due to the activity of separate internal and external NADH dehydrogenases and not the result of physically compromised mitochondria. Furthermore, the near-complete inhibition of oxygen utilization by the cytochrome c oxidase inhibitor cyanide (~92 and ~94% effective in the presence of endogenous and exogenous NADH, respectively) strongly suggests that oxygen consumption was dependent due to membrane-bound oxidative phosphorylation. In contrast to CBS11895, mitochondria from strains IMX2017 and IMX2197 both exhibited 88% lower oxygen consumption rates in the presence of exogenous NADH, suggesting that mitochondria from these strains do not possess external NADH dehydrogenase activity.).

When assayed with endogenous NADH, mitochondria from strains IMX2017 (‘Complex I only’) and IMX2197 (‘Ndh2-1 only’) exhibited similar coupled oxygen-uptake rates to mitochondria isolated from the wild-type strain CBS11895 (Table 2). Presence of the specific Complex I inhibitor rotenone strongly decreased oxygen-uptake rates with endogenous NADH of mitochondria from the strains CBS11895 and IMX2017 (by 80 and 71%, respectively), while for strain IMX2197 no significant rotenone inhibition was observed. These observations demonstrate that Ndh2-1 is indeed internally oriented and able to completely take over the role of Complex I in NADH oxidation in *Δnubm* mutants under aerobic, glucose-limited conditions.

When mitochondria isolated from the wild-type strain *O. parapolymorpha* CBS11895 were assayed with exogenous NADPH, oxygen consumption was detected at a rate of 0.09 ± 0.01 μmol O_2_ (mg protein)^−1^ min^−1^, substantially lower than the rate obtained with exogenous NADH (0.33 ± 0.05 μmol O_2_ (mg protein)^−1^ min^−1^, Table 2). NADPH-dependent oxygen consumption was not significantly coupled (RCR of 1.1). However, it was largely inhibited by cyanide (~85%), indicating that NADPH oxidation did not occur *via* a soluble enzyme but by an external NADPH-accepting dehydrogenase that transferred electrons from NADPH into the mitochondrial respiratory chain. In strains IMX2017 and IMX2197, exogenous NADPH oxidation activity was reduced by 3- and 9-fold, respectively, indicating the NADPH oxidation observed in CBS11895 is caused by Ndh2-2 and/or Ndh2-3.

### *O. parapolymorpha* mutants with linearized respiratory chains exhibit respiratory physiology in glucose-limited chemostat cultures

In aerobic glucose-limited chemostat cultures grown at a dilution rate of 0.1 h^−1^, both strain IMX2017 *(NUBM Δndh2-1 Δndh2-2 Δndh2-3)* and strain IMX2197 *(Δnubm NDH2-1 Δndh2-2 Δndh2-3)* exhibited essentially the same respiratory physiology as the wild-type strain CBS11895. Despite the deletion of genes encoding respiration-linked NAD(P)H dehydrogenases, both mutant strains grew without detectable formation of fermentation products. Moreover, an oxidative respiratory quotient close to 1 was observed and the carbon contained in the glucose feed could be completely recovered as biomass and carbon dioxide (Table 3). IMX2017 exhibited a biomass yield of 0.52 g biomass (g glucose)^−1^, which is not significantly different from that found with wild type CBS11895 (0.51 g biomass (g glucose)^−1^). In contrast, the biomass yield of strain IMX2197 was reduced by ~15% compared to that of the two other strains. This reduced biomass yield is consistent with a less efficient oxidative respiratory chain, using internal, non-proton pumping Ndh2-1 instead of the proton-pumping Complex I. Accordingly, IMX2197 exhibited an approximately ~30% lower biomass yield on oxygen, with correspondingly higher biomass-specific rates of oxygen consumption and carbon-dioxide production.

### The *O. parapolymorpha* NDH2s oxidize NADH but not NADPH when expressed in *E. coli* membranes

The oxygen-consumption experiments indicated that mitochondria from *O. parapolymorpha* poorly couple oxidation of exogenous NADPH to oxygen consumption *via* the aerobic respiratory chain. In principle, any external NDH2 could be responsible for this activity. Upon closer examination of the putative amino acid sequence of *O. parapolymorpha* Ndh2-3, an uncharged residue (Q365) instead of a negatively charged (E272 in *S. cerevisiae* Ndi1) is present within the substrate-binding domain (Figure S1). This residue has been suggested to be involved in determining NADH/NADPH specificity due to interaction with the phosphate group of NADPH (13, 14).

To assess the ability of the *O. parapolymorpha* NDH2s to catalyze the oxidation of NADH and/or NADPH, Ndh2-1, Ndh2-2 and Ndh2-3 were individually overexpressed in *Escherichia coli*, following a strategy previously applied to an NDH2 (Ndi1) from *S. cerevisiae* (27). Since respiration-linked NADPH-dehydrogenase activity has not been reported for *E. coli*, this host was regarded as especially suitable to assess NADPH-dehydrogenase activity of heterologously expressed enzymes.

In spectrophotometric assays at pH 7.4, expression of each of the three *O. parapolymorpha* NDH2s led to 2.4 to 3.2-fold higher NADH-oxidation rates than observed with membranes isolated from the parental *E. coli* strain (Figure 1). This activity indicated successful overexpression and localization to the *E. coli* membrane, and confirmed that the three *O. parapolymorpha* NDH2s are indeed NADH:quinone oxidoreductases. When NADPH was added as substrate to the same membrane preparations, no detectable NADH-oxidation was measurable, indicating that none of the three *O. parapolymorpha* NDH2s can effectively utilize NADPH under the conditions tested (Table S1). Since NAD(P)H oxidation by NDH2s can depend on pH (12, 28), NADH and NADPH oxidation measurements were repeated at pH 5.5 and 8.0. These different pH values did not influence NADH oxidation rates relative to rates measured at pH 7.4 (Student’s t test, p >0.05), and did not stimulate NADPH utilization by either endogenous *E. coli* respiratory enzymes or *O. parapolymorpha* NDH2s (Table S1). Finally, oxidative NAD(P)H catalysis by Ndh2-3 overexpressed in *E. coli* was also tested in the presence of calcium, as Ndh2-3 contains a putative EF-hand suggesting a calcium binding domain which could potentially regulate catalytic behavior (Figure S2) (12). However, the presence of 5 mM Ca^2+^ did not significantly affect rates of NADH oxidation by Ndh2-3 overexpression membranes at pH 5.5 and 7.4 (24% reduction at pH 8.0) and did not enable NADPH utilization at pH 5.5, 7.4 or 8.0 (Table S1).

**Figure 1:**
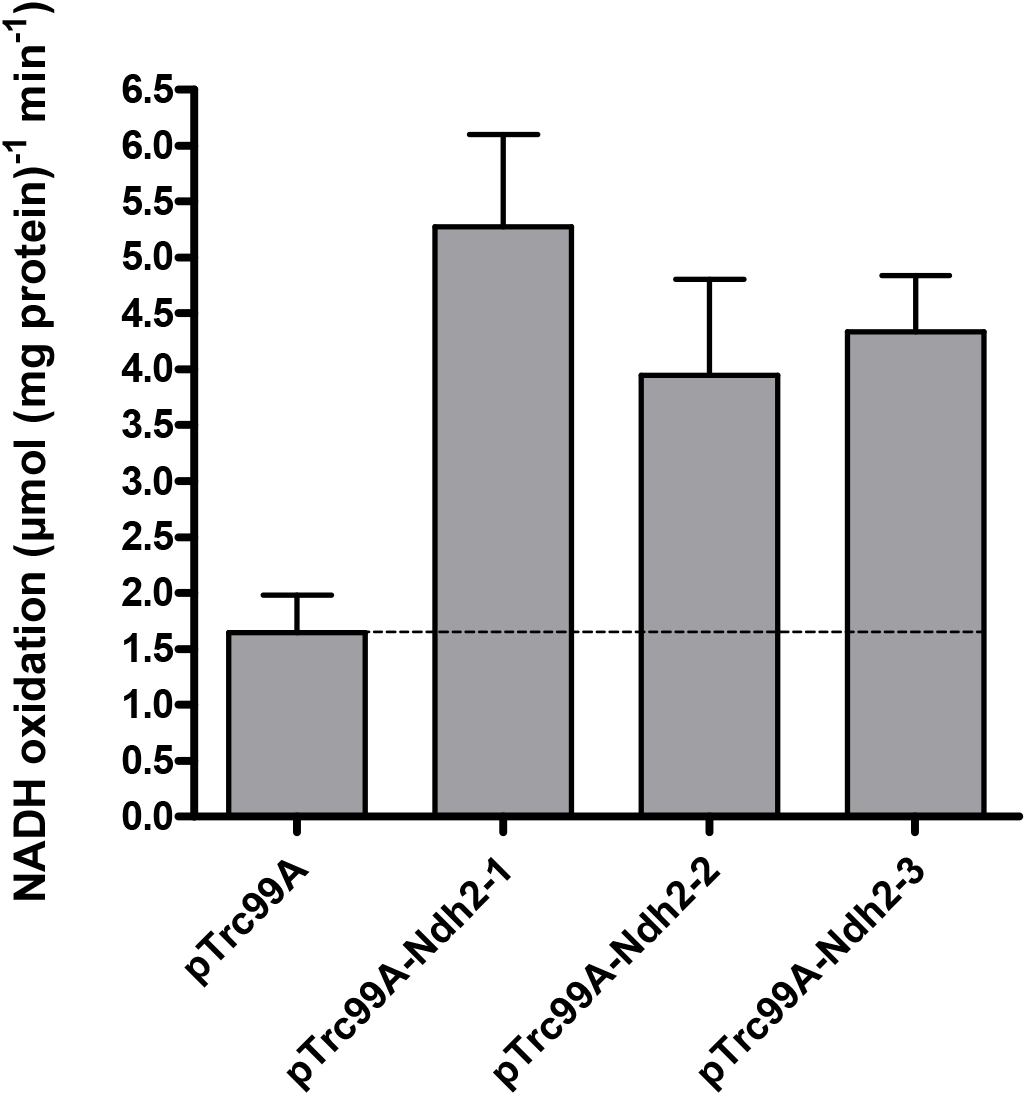
NADH oxidation by *E. coli* membranes isolated from strains overexpressing individual *O. parapolymorpha NDH2s*. Control measurements were done with membranes isolated from a strain carrying an empty overexpression plasmid (pTrc99A). Assays were performed with a membrane protein concentration of 10 μg mL^−1^ at 37°C, with 200 μM NADH and 100 μM ubiquinone-1. Data is presented as mean ± standard deviation of triplicate measurements.

Taken together, these data conclusively show that when assessed in *E. coli* membranes, the three NDH2s from *O. parapolymorpha* can efficiently oxidize NADH at a range of pH levels (pH 5.5-8.0), but do not catalyze the oxidation of NADPH.

## Discussion

### NAD(P)H dehydrogenases in the respiratory chain of *O. parapolymorpha*

In this study, the *O. parapolymorpha* gene HPODL_02792 *(NDH2-1)* was demonstrated to encode a mitochondrial ‘internal’ alternative NADH dehydrogenase. To achieve consistent nomenclature with other fungal species, we suggest naming this gene *OpNDI1* (<u>N</u>ADH <u>d</u>ehydrogenase, <u>i</u>nternal). The closely related yeast *O. polymorpha* likely also possesses an internal alternative NADH dehydrogenase, as its genome (29) encodes a protein (OGAPODRAFT_15258) that exhibits 98% sequence identity with OpNdi1.

Mitochondria isolated from *O. parapolymorpha* also oxidized externally supplied NADH, indicating that either one or both of the remaining alternative dehydrogenases, Ndh2-2 and Ndh2-3, are facing the intermembrane space, rather than the mitochondrial matrix. Coexistence of more than one internal alternative dehydrogenase has been described (30), but since the fungi and yeasts so far characterized either possess one or no such enzymes, both Ndh2-2 and Ndh2-3 appear to be externally oriented. The internal orientation of Ndh2-1/OpNdh1 has now been established in this study, oxygen uptake experiments with mitochondria isolated from strains IMD004 *(Δndh2-2)* and IMD005 *(Δndh2-3)* could be performed to confirm the localization of the other two enzymes.

Wild-type *O. parapolymorpha* mitochondria also oxidized NADPH *via* the respiratory chain as demonstrated by near-complete inhibition of this activity by cyanide. While growth on glucose the production of NADPH can be balanced with biosynthetic requirements, assimilation of other carbon sources such as gluconate result in a surplus of NADPH in yeast (31), making the ability to respire NADPH a beneficial trait. The oxygen-uptake activity with NADPH was essentially uncoupled (RCR of 1.1), which was unexpected, as addition of ADP should relieve the backpressure of the proton gradient, especially since the same mitochondrial preparations exhibited well-coupled oxygen uptake in the presence of NADH. Mitochondria isolated from glucose-limited chemostat cultures of *K. lactis* (D = 0.1 h^−1^) similarly exhibited uncoupled oxygen uptake in the presence of NADPH (32), while mitochondria from lactate- and glucose-grown batch cultures of the same yeast exhibited RCRs of 2.3 and 1.3 for NADPH, respectively (9). With mitochondria isolated from glucose-limited chemostats of *Candida utilis*, the RCRs of NADPH oxidation were found to vary as a function of the dilution rate, with cultures grown at D = 0.05-0.1 h^−1^ exhibiting an RCR of ~1.9 while cultures grown at D = 0.2-0.4 h^−1^ exhibited a lower RCR of ~1.2 (33). In agreement with our study, these studies reported inhibition of NADPH oxidation by cyanide and/or antimycin A, indicating that the degree of oxidative coupling of NADPH respiration by yeast mitochondria is dependent on the utilized substrate and/or the growth condition employed, which might also be the case in *O. parapolymorpha*.

The heterologous expression of *O. parapolymorpha* NDH2s in *E. coli* membranes showed that NADPH oxidation of the *O. parapolymorpha* mitochondria is not likely to have occurred *via* these NDH2(s), since the three enzymes exclusively utilized NADH when assayed within *E. coli* membranes. Based on their protein sequences, we originally speculated Ndh2-3 to be the most likely candidate to accept NADPH, as it contains the uncharged residue proposed to permit NADPH utilization (14). There are indeed NADPH-utilizing NDH2s with this uncharged residue such as Nde1 from *N. crassa* (Figure S1) or plant enzyme St-NBD1 (14), and mutation of this exact residue has been exploited to alter substrate specificity from NADH to co-utilization of NADH and NADPH in a bacterial NDH2 (34). However, Nde2 from *N. crassa* and Nde1 from *K. lactis* do not contain the uncharged residue but still accept both substrates (Figure S1), indicating that a charged amino acid in this position does not strictly prevent NADPH utilization and that NDH2 substrate specificity cannot be accurately predicted by a single residue. In contrast to Nde1 from *N. crassa*, where NADPH oxidation is highly affected by calcium (12), the putative calcium binding domains of Ndh2-3 from *O. parapolymorpha* and Nde2 from *K. lactis* are poorly conserved, and apparently lost their regulatory function, as presence of calcium did not appear to have an effect on activity of either enzyme (9).

### NDH2-independent mechanisms for respiration of cytosolic NADH

Mitochondria from strains IMX2017 *(Δndh2-1 Δndh2-2 Δndh2-3)* and IMX2197 *(Δnubm Δndh2-2 Δndh2-3)* did not oxidize external NADH, but both strains exhibited a fully respiratory phenotype in glucose-limited chemostat cultures at D = 0.1 h^−1^. Since both strains only contain a NADH dehydrogenase that can accept electrons from NADH in the mitochondrial matrix (Complex I and Ndh2-1/OpNdi1, respectively), mechanisms other than the external alternative NADH dehydrogenases are capable of respiring cytosolic NADH in these strains must be present. One candidate would be the ‘Gut2/Gpd shuttle’, consisting of mitochondrial glycerol-3-phosphate dehydrogenase (Gut2), which indirectly respires cytosolic NADH in combination with cytosolic NAD-dependent glycerol 3-phosphate dehydrogenase (Gpd) (36). When grown on glycerol, *O. parapolymorpha* exhibits highly increased glycerol-kinase activity, indicating that glycerol assimilation occurs *via* the phosphorylative pathway. This indicates that Gut2 is functional in this yeast (37). Furthermore, Gut2 activity was demonstrated in the closely related yeast *O. polymorpha* (38). In *S. cerevisiae*, the metabolic function of Gut2 overlaps with the external NDH2s (39, 40), and yeast mitochondria isolated from glucose-grown cultures of various yeast species have been demonstrated to respire glycerol-3-phosphate with similar specific oxygen uptake rates as NADH (7, 22, 32).

While *S. cerevisiae* lacking Gut2 in addition to its two external NADH dehydrogenases produces large amounts of glycerol under aerobic, glucose-limited conditions (40), an *O. parapolymorpha* mutant lacking Gut2 in addition to the three NDH2s *(Δndh2-1 Δndh2-2 Δndh2-3 Δgut2)* exhibited fully respiratory physiology under identical conditions (Table S2). The absence of byproduct formation indicates that at least one additional, unknown mechanism for coupling the oxidation of cytosolic NADH to mitochondrial respiration is present in *O. parapolymorpha*. Such a mechanism could, for example, involve a shuttle for import of NADH equivalents into the mitochondrial matrix, where oxidation by Complex I and/or OpNdi1 can occur. For *S. cerevisiae*, several suitable mechanisms have been demonstrated or proposed, such as the malate-oxaloacetate and malate-aspartate shuttles, mitochondrial oxidation of ethanol produced in the cytosol, or, assuming a high cytosolic NADH/NAD^+^ ratio, an inward-directed ethanol-acetaldehyde shuttle (36, 41). Based on sequence homology with *S. cerevisiae*, key proteins for these mechanisms are present in *O. parapolymorpha* (Table S3), however their operation and physiological relevance has not been confirmed by functional analysis studies.

### Physiological relevance of branched respiratory chains

Respiratory chains of plants, fungi and some protists are branched (3, 42), and, as also demonstrated in this study for *O. parapolymorpha*, can exhibit a metabolic redundancy for respiration-linked NADH oxidation, minimizing the physiological effect of loss of individual or multiple NADH dehydrogenases (10, 40, 43). In general, this redundancy and the exact physiological role of the alternative NADH dehydrogenases are still poorly understood. In *O. parapolymorpha*, OpNdi1 appears to have a unique metabolic function as it is the only NADH dehydrogenase strictly required to sustain fast respiratory growth under glucose excess conditions, demonstrating that limiting respiratory capacity by disruption of a single NADH dehydrogenase can elicit the Crabtree effect in a Crabtree-negative yeast. Similarly, the overexpression of a single ‘Gal4-like’ transcription factor has been reported to convert Crabtree-negative *P. pastoris* into a Crabtree-positive yeast (44). However, since formation of ethanol and CO_2_ from glucose by alcoholic fermentation is redox-neutral and reduced by-products such as glycerol were not detected in fermenting cultures of IMD003 *(Δndh2-1)* and IMX2017 *(Δndh2-1 Δndh2-2 Δndh2-3)*, (re)oxidation of mitochondrial NADH from oxidative sugar metabolism must still occur via respiration. While Complex I has been demonstrated to be physiologically irrelevant under these conditions in wild-type *O. parapolymorpha* CBS11895 (18), based on the available data its contribution to oxidation of mitochondrial NADH cannot be excluded in these strains. Alternatively, if export of NADH equivalents from the mitochondrial matrix is possible in this yeast, NADH could also be oxidized in the cytosol, for example by external NADH dehydrogenase(s) in IMD003 or the Gut2/Gpd shuttle in IMX2017.

A previous study with submitochondrial particles harvested from stationary-phase cultures of *O. polymorpha* found that most NADH oxidation occurred via NDH2 and only ~10% via Complex I (45). Similarly, in our experiments with mitochondria isolated from wild-type *O. parapolymorpha* grown in glucose-limited chemostat cultures, only ~30% of the total specific NADH oxidation activity could be attributed to Complex I. However, due to the saturating substrate concentrations used in these types of experiments, they allow only for a limited interpretation of the actual physiological relevance of the respective enzymes. *In vivo*, competition of Complex I and OpNdi1 for NADH likely occurs if both systems are expressed at the same time, and indeed some fungal species appear to co-utilize Complex I and internal NDH2 (21, 46, 47). With mitochondria isolated from wild-type *O. parapolymorpha* strain CBS11895, the presence of 50 μM rotenone did not fully inhibit internal NADH oxidation. However, as strain IMX2017, which lacked all NADH dehydrogenases besides Complex I, exhibited a similar partial inhibition of oxygen uptake by rotenone, the uninhibited activity in CBS11895 was likely not caused by OpNdi1 but instead by incomplete inhibition of *O. parapolymorpha* Complex I. Comparative studies with submitochondrial particles have demonstrated that rotenone inhibits Complex I from yeasts less strongly than the Bos taurus enzyme, requiring 50 μM rotenone to achieve 96% inhibition of NADH oxidation activity by the *P. pastoris* Complex I (45). It is conceivable that Complex I from *O. parapolymorpha* is even more resistant to rotenone, explaining the observed partial inhibition in strains CBS11895 and IMX2017. In addition, OpNdi1 was not detected in the proteome of aerobic, glucose-limited cultures of CBS11895 (18), and the observed identical biomass yields of strains CBS11895 and IMX2017 are consistent with a situation in which essentially all mitochondrial NADH is (re)oxidized by Complex I in wild-type *O. parapolymorpha* under these conditions. These observations indicate that oxidation of mitochondrial NADH is strictly separated between Complex I and OpNdi1 in *O. parapolymorpha* under conditions of glucose limitation and glucose excess, respectively. Nevertheless, OpNdi1 is able to fully support respiratory growth in the absence of a functional Complex I, as previously hypothesized (18).

### Concluding remarks

In this study we show that the Crabtree-negative yeast *O. parapolymorpha* contains both NDH1- and NDH2-type NADH dehydrogenases for respiration of NADH in the mitochondrial matrix, but limits their utilization to conditions of carbon limitation and carbon excess, respectively. Furthermore, we find that the respiratory chain of *O. parapolymorpha* can tolerate multiple deletions without compromising respiratory metabolism, offering insight into its flexible nature and opportunities for metabolic (redox) engineering in this industrially-relevant yeast. Finally, the phenotype elicited by disruption of OpNdi1 demonstrates that limiting respiratory capacity by a single mutation in an NADH dehydrogenase can result in overflow metabolism and convert *O. parapolymorpha* into a yeast with a Crabtree-positive phenotype.

## Materials and Methods

### Yeast strains and maintenance

The *O. parapolymorpha* strains used in this study are derived from the wild-type strain CBS11895 (DL-1; ATCC26012) and are described in Table 4. For construction and maintenance, yeast strains were grown in an Innova shaker incubator (New Brunswick Scientific, Edison, NJ, USA) set to 30°C and 200 rpm, in 500 mL shake flasks containing 100 mL heat-sterilized (120°C for 20 min) YPD medium (10 g L^−1^ Bacto yeast extract, 20 g L^−1^ Bacto peptone, 20 g L^−1^ glucose, demineralized water). Solid medium was prepared by addition of 2% (w/v) agar. Frozen stock cultures were prepared from exponentially growing shake-flask cultures by addition of glycerol to a final concentration of 30% (v/v), and aseptically stored in 1 mL aliquots at −80°C.

**Table 4:**
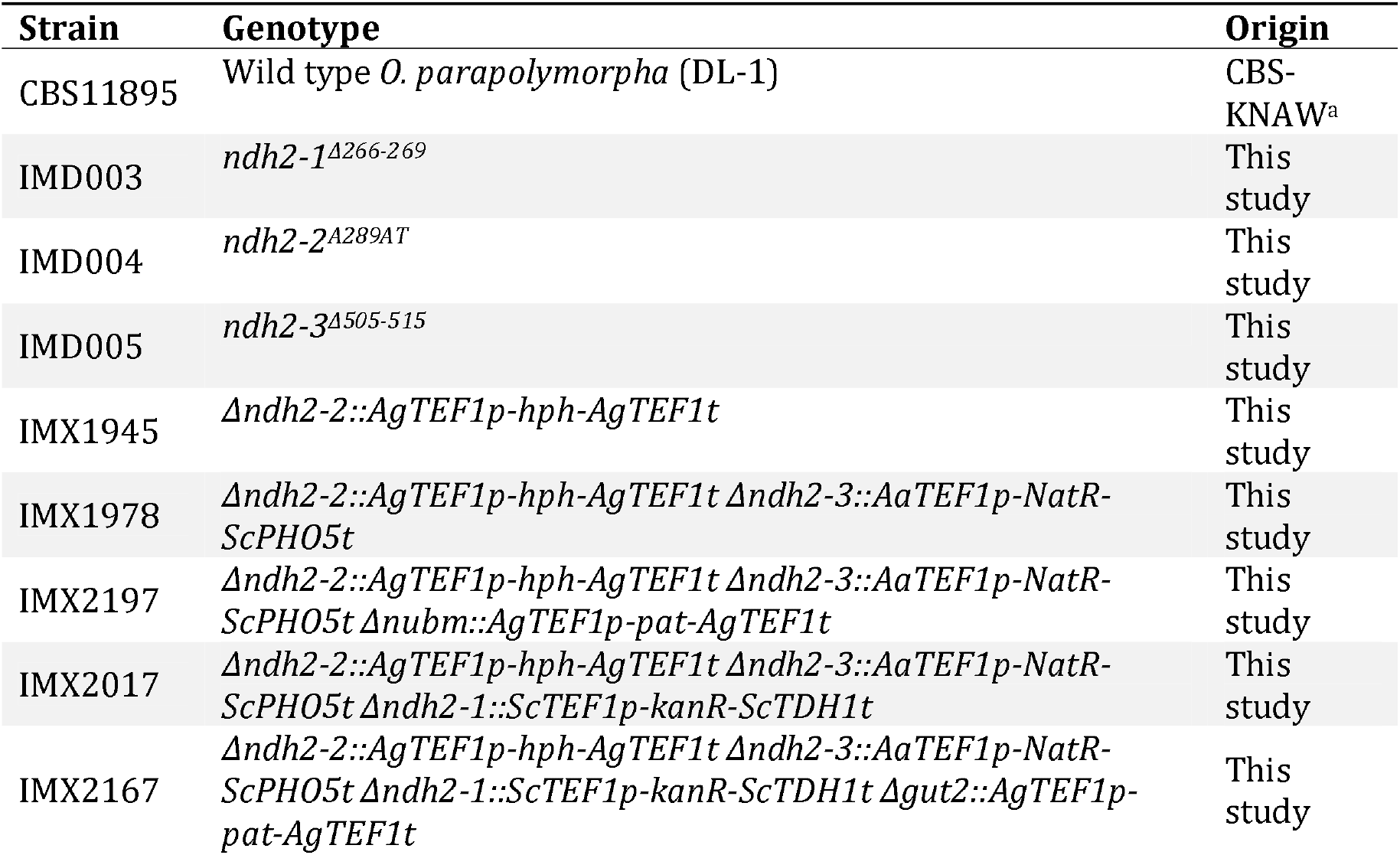
*O. parapolymorpha (Hansenula polymorpha)* strains used in this study. ^a^CBS11895 was obtained from the CBS-KNAW fungal collection (Westerdijk Fungal Biodiversity Institute, Utrecht, The Netherlands).

| Strain | Genotype | Origin |
| --- | --- | --- |
| CBS11895 | Wild type *O. parapolymorpha* (DL-1) | CBS-KNAW<sup>a</sup> |
| IMD003 | *ndh2-1*<sup>Δ266-269</sup> | This study |
| IMD004 | *ndh2-2*<sup>Δ289AT</sup> | This study |
| IMD005 | *ndh2-3*<sup>Δ505-515</sup> | This study |
| IMX1945 | *Δndh2-2::AgTEF1p-hph-AgTEF1t* | This study |
| IMX1978 | *Δndh2-2::AgTEF1p-hph-AgTEF1t Δndh2-3::AaTEF1p-NatR-ScPHO5t* | This study |
| IMX2197 | *Δndh2-2::AgTEF1p-hph-AgTEF1t Δndh2-3::AaTEF1p-NatR-ScPHO5t Δnubm::AgTEF1p-pat-AgTEF1t* | This study |
| IMX2017 | *Δndh2-2::AgTEF1p-hph-AgTEF1t Δndh2-3::AaTEF1p-NatR-ScPHO5t Δndh2-1::ScTEF1p-kanR-ScTDH1t* | This study |
| IMX2167 | *Δndh2-2::AgTEF1p-hph-AgTEF1t Δndh2-3::AaTEF1p-NatR-ScPHO5t Δndh2-1::ScTEF1p-kanR-ScTDH1t Δgut2::AgTEF1p-pat-AgTEF1t* | This study |

### Plasmid construction

All plasmids used in this study are described in Table 5. Plasmids pUD546, pUD547 and pUD548 were *de novo* synthesized by GeneArt (Thermo Fisher Scientific, Waltham, MA, USA) and contained synthetic guide RNA (gRNA) constructs with spacer sequences (5’-3’) ‘CCTGATGTAAATATACGCTG’, ‘AAGAAGAACATTGTTATTCT’, and ‘GTTATTCTGGGTTCCGGCTG’, respectively. Cas9/gRNA co-expression plasmids pUDP019, pUDP020 and pUDP021 were constructed using pUD546, pUD547 and pUD548, respectively, by integration into pUDP002 via BsaI-mediated ‘Golden Gate’ assembly (48) as described previously (49), and verified by digestion with PdmI.

**Table 5:** Plasmids used in this study. Restriction enzyme sites are indicated in superscript, and genes targeted by gRNAs or homologous recombination are indicated in subscript. Aa: *Arxula adeninivorans*; Sp: *Streptococcus pyogenes*; Ag: *Ashbya gossypii*; Sc: *Saccharomyces cerevisiae*; HH: hammerhead ribozyme; HDV: hepatitis delta virus ribozyme; MCS: multiple cloning site; HRS: homologous recombination sequence.

| Name | Relevant characteristics | Origin |
| --- | --- | --- |
| pUD546 | BsaI<sup>HH-gRNA<sub>NDH2-1</sub></sup>-HDV<sup>BsaI</sup> | GeneArt |
| pUD547 | BsaI<sup>HH-gRNA<sub>NDH2-2</sub></sup>-HDV<sup>BsaI</sup> | GeneArt |
| pUD548 | BsaI<sup>HH-gRNA<sub>NDH2-3</sub></sup>-HDV<sup>BsaI</sup> | GeneArt |
| pUDP002 | *panARS (OPT) AgTEF1p-hph-AgTEF1t ScTDH3p<sup>BsaI BsaI</sup>ScCYC1t AaTEF1p-Spcas9 -ScPHO5t* | (49) |
| pUDP019 | *panARS (OPT) AgTEF1p-hph-AgTEF1t ScTDH3p-HH-gRNA<sub>NDH2-1</sub>-HDV-ScCYC1t AaTEF1p-Spcas9 -ScPHO5t* | This study |
| pUDP020 | *panARS (OPT) AgTEF1p-hph-AgTEF1t ScTDH3p-HH-gRNA<sub>NDH2-2</sub>-HDV-ScCYC1t AaTEF1p-Spcas9 -ScPHO5t* | This study |
| pUDP021 | *panARS (OPT) AgTEF1p-hph-AgTEF1t ScTDH3p-HH-gRNA<sub>NDH2-3</sub>-HDV-ScCYC1t AaTEF1p-Spcas9 -ScPHO5t* | This study |
| pUD602 | Template for plasmid backbone (*ori ampR*) | (49) |
| pYTK013 | Template for *ScTEF1p* promoter | (70) |
| pYTK056 | Template for *ScTDH1t* terminator | (70) |
| pYTK077 | Template for *kanR* ORF | (70) |
| pYTK078 | Template for *NatR* ORF | (70) |
| pAG31 | Template for *AgTEF1p-pat-AgTEF1t* cassette | (71) |
| pUD740 | Template for *ScTEF1p-kanR-ScTDH1t* cassette | This study |
| pUD801 | HRS<sub>NDH2-1</sub>-*ScTEF1p-kanR-ScTDH1t*-HRS<sub>NDH2-1</sub> | This study |
| pUD802 | HRS<sub>NDH2-2</sub>-*AgTEF1p-hph-AgTEF1t*-HRS<sub>NDH2-2</sub> | This study |
| pUD803 | HRS<sub>NDH2-3</sub>-*AaTEF1p-NatR-ScPHO5t*-HRS<sub>NDH2-3</sub> | This study |
| pUD1035 | HRS<sub>GUT2</sub>-*AgTEF1p-pat-AgTEF1t*-HRS<sub>GUT2</sub> | This study |
| pUD1036 | HRS<sub>NUBM</sub>-*AgTEF1p-pat-AgTEF1t*-HRS<sub>NUBM</sub> | This study |
| pTrc99A | *ori ampR trp/lac-MCS-rrnB* | (72) |
| pTrc99A-NDH2-1 | *ori ampR trp/lac-NDH2-1-rrnB* | This study |
| pTrc99A-NDH2-2 | *ori ampR trp/lac-NDH2-2-rrnB* | This study |
| pTrc99A-NDH2-3 | *ori ampR trp/lac-NDH2-3-rrnB* | This study |

Construction of pUD740 was done via ‘Gibson assembly’ (50) from the following PCR-amplified fragments: the *ScTEF1* promoter from pYTK013 (primers 12099 + 12100), the *kanR* G418 resistance marker ORF from pYTK077 (primers 12097 + 12098), the *ScTDH1* terminator from pYTK056 (primers 12095 + 12096), the *AaTEF1p-Spcas9-ScPHO5t* expression cassette from pUDP002 (primers 10426 + 10427), upstream (primers 12093 + 12094) and downstream (primers 12101 + 12102) homologous recombination sequences of the *OpKU70* locus from CBS11895 genomic DNA and the plasmid backbone from pUD602 (primers 12103 + 12104).

Plasmids pUD801, pUD802, pUD803, pUD1035 and pUD1036 carrying subcloned homology-flanked marker cassettes for split-marker deletion were constructed by Gibson assembly from various PCR-amplified fragments. For the construction of pUD801 these fragments were: the *ScTEF1p-kanR-ScTDH1t* G418 resistance marker cassette from pUD740 (primers 12100 + 4377), upstream (primers 14385 + 14386) and downstream (primers 14387 + 14388) homologous recombination sequences of the *NDH2-1* locus from CBS11895 genomic DNA and the plasmid backbone from pUD602 (primers 12103 + 12104). For pUD802 these fragments were: the *AgTEF1p-hph-AgTEF1t* hygromycin resistance marker cassette from pUDP002 (primers 11065 + 11133), upstream (primers 14389 + 14390) and downstream (primers 14391 + 14392) homologous recombination sequences of the *NDH2-2* locus from CBS11895 genomic DNA and the plasmid backbone from pUD602 (primers 12103 + 12104). For pUD803 these fragments were: the *AaTEF1* promoter from pUDP002 (primers 10426 + 14383), the *NatR* nourseothricin resistance marker ORF from pYTK078 (primers 14381 + 14382), the *ScPHO5* terminator from pUDP002 (primers 10427 + 14384), upstream (primers 14393 + 14394) and downstream (primers 14395 + 14396) homologous recombination sequences of the *NDH2-3* locus from CBS11895 genomic DNA and the plasmid backbone from pUD602 (primers 12103 + 12104). For pUD1035 these fragments were: the *AgTEF1p-pat-AgTEF1t* phosphinothricin resistance marker cassette from pAG31 (primers 3242 + 8439), upstream (primers 14803 + 14804) and downstream (primers 14805 + 14806) homologous recombination sequences for *GUT2* from CBS11895 genomic DNA and the plasmid backbone from pUD602 (primers 12103 + 12104). For pUD1036 these fragments were: the *AgTEF1p-pat-AgTEF1t* phosphinothricin resistance marker cassette from pAG31 (primers 3242 + 8439), upstream (primers 14807 + 14808) and downstream (primers 14809 + 14810) homologous recombination sequences for *NUBM* from CBS11895 genomic DNA and the plasmid backbone from pUD602 (primers 12103 + 12104). Correct insertion and presence of the homology-flanked marker cassettes in the constructed plasmids was verified by restriction digest and diagnostic PCR using PvuI + NdeI and primer sets 2908 + 12616 and 1642 + 3983 (pUD801), PvuI + PsiI and primer sets 2457 + 12616 and 1642 + 1781 (pUD802), PvuI + KpnI and primer sets 10459 + 12616 and 1642 + 10458 (pUD803) and PvuI + NcoI and primer sets 1409 + 12616 and 1642 + 4662 (pUD1035 & pUD1036).

Plasmids pTrc99A-NDH2-1, pTrc99A-NDH2-2 and pTrc99A-NDH2-3 were constructed by restriction/ligation cloning. The ORFs encoding *NDH2-1* (HPODL_02792), *NDH2-2* (HPODL_00256) and *NDH2-3* (HPODL_02018) were PCR-amplified from CBS11895 genomic DNA using primer sets 14929 + 14931, 16075 + 16076 and 16077 + 16078, respectively. Amplification using these primer sets added a 5’ NcoI site, a GS-flanked 6-HIS tag (‘GSHHHHHHGS’) directly after the start codon and a 3’ XmaI site to all three ORFs. Furthermore, amplification of *NDH2-1* omitted the first 24 amino acids after the start codon, as they could be unambiguously identified as mitochondrial targeting sequence by MitoFates (51). PCR amplicons were then digested with NcoI and XmaI and cloned into NcoI/XmaI-digested pTrc99A using T4 DNA ligase (New England Biolabs, Ipswich, MA, USA).

### Yeast strain construction

*O. parapolymorpha* strains were transformed via electroporation of freshly prepared electrocompetent cells as described previously (49). Depending on the selection marker, mutants were selected on solid YPD medium supplemented with 200 μg mL^−1^ G418, 300 μg mL^−1^ hygromycin B or 100 μg mL^−1^ nourseothricin, or on solid synthetic medium (SM) supplemented with 20 g L^−1^ glucose and 200 μg mL^−1^ bialaphos (SanBio, Uden, The Netherlands). SM was prepared according to Verduyn, Postma (52) and autoclaved at 120°C for 20 min. Glucose and vitamins (52) were prepared separately and filter-sterilized (vitamins) or heat-sterilized at 110°C for 20 min (glucose).

Strains IMD003, IMD004 and IMD005 with disrupted versions of genes *NDH2-1* (HPODL_02792), *NDH2-2* (HPODL_00256) and *NDH2-3* (HPODL_02018), respectively, were constructed using the pUDP CRISPR/Cas9 system described previously (49). Wild type strain CBS11895 was transformed with pUDP019, pUDP020 or pUDP021 targeting *NDH2-1* (after base part 269 out of 1614), *NDH2-2* (after base pair 290 out of 1671) and *NDH2-3* (after base pair 515 out of 2097), respectively, and subjected to the prolonged liquid incubation protocol as described previously for deletion of *OpADE2* and *OpKU80* (49). Randomly picked colonies were then subjected to PCR amplification of the *NDH2-1, NDH2-2* and *NDH2-3* locus using primer sets 10742 + 10743, 10744 + 10745 and 10746 + 10747, respectively, followed by Sanger sequencing (Baseclear, Leiden, The Netherlands) to identify mutant transformants harboring a frame-shifting indel at the respective gRNA target sites. Three mutants with either a deletion of base pairs 226-229 of *NDH2-1*, an additional thymine nucleotide between position 289 and 290 of *NDH2-2* or a deletion of base pairs 505-515 of *NDH2-3* were identified, restreaked three times subsequently on non-selective YPD medium to remove the pUDP plasmids, and renamed IMD003, IMD004 and IMD005, respectively.

Strains IMX1945, IMX1978, IMX2017, IMX2197 and IMX2167 were constructed using a split-marker deletion approach (53), with ~480 bp of internal (marker recombination) and ~480 bp of external (genome recombination) homology. To preserve the promoter and terminator sequences of neighboring genes and limit interference of their expression, a minimum of 800 bp preceding and 300 bp succeeding adjacent ORFs were kept unaffected by the deletions. IMX1945 was constructed from wild type *O. parapolymorpha* strain CBS11895 by deletion of *NDH2-2* with a *hyg* resistance cassette using pUD802, IMX1978 was constructed from IMX1945 by additional deletion of *NDH2-3* with a *NatR* resistance cassette using pUD803, IMX2017 was constructed from IMX1978 by additional deletion of *NDH2-1* with a *kanR* resistance cassette using pUD801, IMX2197 was constructed from IMX1978 by additional deletion of *NUBM* (HPODL_04625) (18) with a *pat* resistance cassette using pUD1036, and IMX2167 was constructed from IMX2017 by additional deletion of *GUT2* (HPODL_00581) with a *pat* resistance cassette using pUD1035. For the split-marker deletion of *NDH2-1, NDH2-2, NDH2-3, NUBM* and *GUT2*, the two overlapping fragments for transformation were PCR-amplified from pUD801 using primer sets 6816 + 14397 and 12565 + 14398, from pUD802 using primer sets 14399 + 14400 and 14401 + 14402, from pUD803 using primer sets 14403 + 14404 and 14405 + 14406, from pUD1036 using primer sets 15884 + 15885 and 15886 + 15887, and from pUD1035 using primer sets 14811 + 15885 and 14812 + 15886, respectively. Prior to transformation, the amplified fragments were gel-purified and, in case DNA amounts were too low for transformation, used as template for another PCR amplification using the same primers followed by PCR purification. For each transformation, a total of ~1 μg purified DNA (both fragments equimolar) in a maximum volume of 4 μL was transformed to 40 μL of fresh electrocompetent cells as described above, with the exception that after electroporation the cell suspensions were recovered in 1 mL YPD for 3 h at 30°C before plating onto selective medium. Additionally, after YPD recovery, cells transformed with the *pat* resistance marker were washed once by centrifugation and resuspension in sterile demineralized water before selective plating. Selection plates were typically incubated for 3 days at 30°C before assessment of the correct replacement of the target genes with the resistance markers via diagnostic PCR using primer sets 14465 + 14466, 14465 + 4047 and 2653 + 14466 for *ΔNDH2-1::kanR*, primer sets 14467 + 14468, 14467 + 7864 and 8411 + 14468 for *ΔNDH2-2::hph*, primer sets 14469 + 14470, 14469 + 11197 and 11202 + 14470 for *ΔNDH2-3::NatR*, primer sets 15630 + 15631, 15630 + 15885 and 15886 + 15631 for *ΔNUBM::pat* and primer sets 15628 + 15629, 15628 + 15885 and 15886 + 15629 for *ΔGUT2::pat*. Single colonies that contained the desired genotype(s) were restreaked once on selective medium, followed by two restreaks on non-selective YPD medium before stocking.

### Molecular biology

PCR amplification for cloning and construction was performed with Phusion High Fidelity Polymerase (Thermo Fisher Scientific) using PAGE-purified oligonucleotide primers (Sigma-Aldrich, St. Louis, MO, USA) according to manufacturer’s recommendations, with the exception that a final primer concentration of 0.2 μM was used. Diagnostic PCR was done using DreamTaq polymerase (Thermo Fisher Scientific) and desalted primers (Sigma-Aldrich). The primers used in this study are shown in Table S4. Genomic DNA of yeast colonies was isolated using the LiAc-sodium dodecyl sulfate method (54) or the YeaStar Genomic DNA kit (Zymo Research, Irvine, CA, USA). DNA fragments obtained by PCR were separated by gel electrophoresis. Gel-purification was carried out using the Zymoclean Gel DNA Recovery Kit (Zymo Research). PCR purification was performed using the GenElute PCR Clean-Up Kit (Sigma-Aldrich). Gibson assembly was done using the NEBuilder HiFi DNA Assembly Master Mix (New England Biolabs) with purified DNA fragments according to manufacturer’s recommendations, with the exception that reaction volume was down-scaled to 5-10 μL. DNA fragments that were PCR-amplified from a template harboring the same bacterial resistance marker as the construct to be Gibson-assembled were subjected to DpnI treatment prior to PCR cleanup. Restriction digest was performed using FastDigest enzymes (Thermo Fisher Scientific) or High Fidelity (HF) restriction endonucleases (New England Biolabs, Ipswich, MA, USA) according to the manufacturer’s instructions. *E. coli* strains XL1-blue and DH5α were used for plasmid transformation, amplification and storage. Plasmid isolation from *E. coli* was done using the GenElute Plasmid Miniprep Kit (Sigma-Aldrich) or the Monarch Plasmid Miniprep Kit (New England Biolabs).

### Multiple sequence alignment and domain prediction

Alignment of NDH2 protein sequences was done using MUSCLE (https://www.ebi.ac.uk/Tools/msa/muscle/) (55) and visualized using Jalview (56). Orientation and substrate specificity of other fungal NDH2s were taken from: *K. lactis* (8, 9), *N. crassa* (10–12, 57), *S. cerevisiae* (6, 7, 58, 59), and *Y. lipolytica* (60). Prediction of the putative EF-hand calcium binding domain in NDH2 sequences was done by Motif Scan (https://myhits.isb-sib.ch/cgi-bin/motif_scan) using PROSITE profiles (61).

### Shake-flask cultivation

Shake-flask growth experiments were performed with synthetic medium with urea as nitrogen source (62), set to an initial pH of 5.0 with KOH. Cultures were grown in 500 mL round-bottom shake flasks filled with 50 mL medium. Cultures were grown with 2 g L^−1^ glucose as sole carbon source and were inoculated with mid-exponential precultures (washed once with sterile demineralized water) to an initial OD_660_ of 0.3. Precultures were grown under the same conditions and in the same medium, but with an initial glucose concentration of 5 g L^−1^. Shake flasks were continuously shaken during sampling to prevent oxygen limitation. Physiological parameters were calculated from at least 5 samples taken during the exponential growth phase. Calculated ethanol yields were not corrected for evaporation.

### Chemostat cultivation

Chemostat cultivation was performed as described previously (18) using SM with the addition of 0.15 g L^−1^ Pluronic 6100 PE antifoaming agent (BASF, Ludwigshafen, Germany) and glucose (7.5 or 9 g L^−1^) as sole carbon source. SM was prepared according to Verduyn, Postma (52) as described above. Bioreactors were inoculated with exponentially growing shake flask cultures (SM with 20 g L^−1^ glucose). Chemostat cultivation was performed in 2-L benchtop bioreactors (Applikon, Delft, The Netherlands) with a working volume of 1.0 L which was maintained by an electrical level sensor that controlled the effluent pump. The dilution rate was set by maintaining a constant medium inflow rate. Cultures were sparged with dried, compressed air (0.5 vvm) and stirred at 800 rpm. Temperature was maintained at 30°C and pH was controlled at 5.0 by automatic addition of a 2 M KOH by an EZcontroller (Applikon). The exhaust gas was cooled with a condenser (2°C) and dried with a Perma Pure Dryer (Inacom Instruments, Veenendaal, the Netherlands) prior to online analysis of carbon dioxide and oxygen with a Rosemount NGA 2000 Analyzer (Emerson, St. Louis, MO, USA). Cultures were assumed to have reached steady state when, after a minimum of 5 volume changes, the oxygen-consumption rate, carbon-dioxide production rate and biomass concentration changed by less than 3% over two consecutive volume changes.

### Analytical methods

Optical density (OD) of yeast cultures was measured at 660 nm on a Jenway 7200 spectrophotometer (Jenway, Staffordshire, UK). OD of bacterial cultures was measured at 600 nm on Ultrospec 2100 pro (Amersham, Little Chalfont, UK). For biomass dry weight determination of yeast cultures, exactly 10 mL of culture broth was filtered over pre-dried and pre-weighed membrane filters (0.45 μm, Pall corporation, Ann Arbor, MI, USA), which were washed with demineralized water, dried in a microwave oven at 350 W for 20 min and weighed immediately (63). Samples were diluted with demineralized water prior to filtration to obtain a biomass dry weight concentration of approximately 2 g L^−1^. The exact dilution was calculated by weighing the amount of sample and diluent and assuming a density of 1 g mL^−1^ for both fractions. Concentrations of extracellular metabolites and putative alcoholic contaminants of NAD(P)H substrates were analyzed by high-performance liquid chromatography (HPLC) on an Agilent 1100 HPLC (Agilent Technologies, Santa Clara, CA USA) with an Aminex HPX-87H ion-exchange column (BioRad, Veenendaal, The Netherlands) operated at 60°C with 5 mM H_2_SO_4_ as mobile phase at a flow rate of 0.6 mL min^−1^. For the determination of extracellular metabolites, 1 mL aliquots of culture broth were centrifuged for 3 min at 20,000 g and the supernatant was used for analysis. Protein concentrations of mitochondrial preparations were estimated by the Lowry method (64), using dried bovine serum albumin (BSA, fatty acid-free, Sigma-Aldrich) as standard. Where necessary, protein determinations were corrected for BSA present in the mitochondrial preparations. Protein concentrations of *E. coli* membrane fractions were determined using a bicinchoninic acid (65) protein assay kit (Sigma-Aldrich) with BSA (Interchim, Montlucon, France) as standard.

### Isolation of mitochondrial fractions

Mitochondria were isolated from glucose-limited, aerobic chemostat cultures (D = 0.1 h^−1^) according to a procedure similar as described for *S. cerevisiae* (7), based on the mild osmotic lysis method developed for *Candida utilis* (66). Biomass (1.5 g dry weight) was harvested by centrifugation at 3000 g for 4 min. The pellet was then resuspended by vortexing in 30 mL of Tris buffer (100 mM) containing 10 mM dithiothreitol (final buffer pH of 9.3) and incubated at 30°C for 10 min. Afterwards, the cells were washed twice with 30 mL buffer A (25 mM potassium phosphate, 2 M sorbitol, 1 mM MgCl2, 1 mM EDTA, pH 7.5) by centrifugation (4000 g, 4 min) and resuspension (gentle vortexing). Then, cells were pelleted again by centrifugation (5000 g, 8 min) and resuspended (gentle vortexing) in a total volume of 40 mL buffer A. 3.06 mg of zymolyase (from *Arthrobacter luteus*, 20,000 U g^−1^, AMS Biotechnology, Abingdon, UK) dissolved in 200 μL buffer A was added to the cell suspension, which was subsequently incubated at 30°C under gentle shaking for 60-90 min. Incubation time depended on the rate of spheroplast formation which was estimated based on the sensitivity to osmotic shock by 200-fold dilution in demineralized water as described previously (66). Incubation was continued until osmotic resistance decreased to approx. 25% (see Figure S3). During the zymolyase treatment, release of glucose-6-phosphate dehydrogenase activity from compromised cells was measured as described previously (66, 67) and typically did not exceed 5% compared to a sonicated sample. After zymolyase treatment, all subsequent steps were carried out on ice or in a cooled (4°C) centrifuge. Spheroplasts were washed twice with 35 mL buffer A by centrifugation (4400 g, 7 min) and resuspension (gentle shaking), followed by centrifugation (5000 g, 6 min) and resuspension (gentle shaking) in a total volume of 10 mL buffer A. Subsequently, 30 mL of buffer B (25 mM potassium phosphate, 0.2 M sorbitol, 1 mM MgCl2, 1 mM EDTA, pH 7.5) was added dropwise to the spheroplast suspension over a timeframe of ~2-3 h, while it was slowly stirred with a magnetic stirrer bar. The spheroplast suspension was then subjected to 2 strokes in a cooled Potter-Elvehjem homogenizer (150 rpm, clearance 28 μm). After centrifugation (3000 g, 10 min), the supernatant was separated from intact cells and debris and spun again (12000 g, 10 min). The resulting pellet, containing the mitochondria, was resuspended in 2.5 mL of buffer C (25 mM potassium phosphate, 0.65 M sorbitol, 1 mM MgCl_2_, 1 mM EDTA, 1 mg mL^−1^ BSA (fatty acid-free, Sigma-Aldrich), pH 7.5) and kept on ice.

### Oxygen-uptake studies with mitochondrial preparations

Substrate-dependent oxygen consumption rates of mitochondria were determined polarographically at 30°C with a Clark-type oxygen electrode. The assay mixture (4 mL) contained 25 mM potassium phosphate buffer (pH 7.0), 5 mM MgCl_2_, and 0.65 M sorbitol. Reactions were started with ethanol (5 mM), methanol (5 mM), L-malate + pyruvate (both 5 mM, adjusted to pH 7.0 with KOH), 0.25 mM NADH (Prozomix, Haltwhistle, UK) or 0.75 mM NADPH (Oriental Yeast Co., Tokyo, Japan). While some commercial preparations of NADH and NADPH are contaminated with ethanol (7, 32, 68), no ethanol (or methanol) was detected via HPLC analysis in freshly prepared, concentrated (100 mM) solutions of the NADH and NADPH used in this study (detection limit: ethanol 1 mM; methanol 5 mM). Oxygen uptake rates were calculated based on a dissolved oxygen concentration of 236 μM in air-saturated water at 30°C. Respiratory control values were determined by adding 0.25 mM ADP (69). For tests with rotenone (50 μM), a concentrated stock solution (20 mM in DMSO) was freshly prepared directly before the respective assays and kept at room temperature. Mitochondria were pre-incubated in the presence of rotenone for 5 min at assay conditions prior to substrate addition. Preincubation with equivalent amounts of DMSO without rotenone did not measurably affect oxygen uptake rates. Tests with KCN (1 mM) were conducted with a concentrated stock solution (200 mM, in 100 mM NaOH), which was added to mitochondria during ADP-stimulated respiration (state III). Addition of equivalent amounts of NaOH without KCN affected oxygen uptake rates by less than 15%.

### Overexpression of *O. parapolymorpha* NDH2s in *E. coli*

Overexpression and purification strategy were based on previous literature (19). *E. coli* BL21(DE3) cells were transformed with plasmids pTrc99A, pTrc99A-NDH2-1, pTrc99A-NDH2-2, and pTrc99A-NDH2-3. Correct transformants were pre-cultured (37°C, 180 rpm) for 16 h in lysogeny broth with 100 μg mL^−1^ ampicillin. These precultures were used to inoculate 500 mL apple flasks with 100 mL 2xYT medium (16 g L^−1^ tryptone, 10 g L^−1^ yeast extract, 5 g L^−1^ NaCl) supplemented with 20 g L^−1^ glucose and 100 μg mL^−1^ ampicillin and grown under the same conditions. Once cultures had reached an OD_600_ of 0.5, NDH2 overexpression was induced using 1 mM isopropyl β-D-1-thiogalactopyranoside (IPTG, Sigma Aldrich), followed by growth for an additional 5 h at 30°C and 180 rpm. Afterwards, cells were harvested by centrifugation (7000 g, 10 min) and washed with 25 mL buffer W1 (50 mM Tris-HCl, pH 8.0, 2 mM MgCl_2_). Cell pellets were then resuspended in buffer W1 (at 4 mL per g wet weight), containing 0.1 mM PMSF (Sigma-Aldrich) and 0.1 mg mL^−1^ bovine pancreatic DNase (New England Biolabs), and incubated 10 min at room temperature. Cells were then disrupted using a cell disruptor (Constant Systems Ltd., Daventry, UK) at 1.38 kbar. Unbroken cells and cell debris were removed by centrifugation (10,000 g, 10 min). The membrane fraction was isolated from the cell lysate by ultracentrifugation (180,000 g, 45 min, 4°C), and the resulting membrane pellet was resuspended in buffer W1 and stored at −80°C.

### *In vitro* NAD(P)H dehydrogenase activity tests with NDH2s

NAD(P)H:quinone oxidoreductase activity was measured using a spectrophotometric assay similar to as described previously (19). Activity was monitored spectrophotometrically using a modified Cary 60 UV/Vis Spectrophotometer (Agilent Technologies), following the oxidation of NADH or NADPH at 340 nm in the presence of ubiquinone-1 (UQ-1, Sigma-Aldrich) at 37°C. Membrane preparations (10 μg protein mL^−1^) and UQ-1 (100 mM) were added to pre-warmed reaction buffer (final volume 2 mL) in a 1 cm path length cuvette and incubated for 30 s. Depending on the pH, the reaction buffer consisted of i) 50 mM Tris-HCl (pH 8.0), 150 mM NaCl, ii) 25 mM MES + 25 mM MOPS (pH 7.4), 150 mM NaCl or iii) 25 mM MES + 25 mM MOPS (pH 5.5), 150 mM NaCl. NADH or NADPH (200 μM at pH 8.0 and 7.4, 100 μM at pH 5.5) were added to the mixture to initiate the reaction. An extinction coefficient of 6.3 mM^−1^ cm^−1^ was used to calculate NAD(P)H concentration. For tests with calcium, 5 mM CaCl_2_ was added to the assay after 1 min of reaction time had elapsed.

## Acknowledgements

The authors thank Veronica Gast for help with plasmid and strain construction, Lisan Broekman for performing part of the shake flask characterization, Marijke Luttik for providing technical expertise and assistance regarding isolation and measurement of mitochondria, Erik de Hulster for assistance with chemostat cultivation and Jack Pronk for helpful discussions and feedback on the manuscript.

This work was performed within the BE-Basic R&D Program (http://www.be-basic.org/), which was granted a FES subsidy from the Dutch Ministry of Economic Affairs, Agriculture and Innovation (EL&I).

## Data availability

All strains and plasmids constructed in this work are available upon request.

---

**Figure S1:**
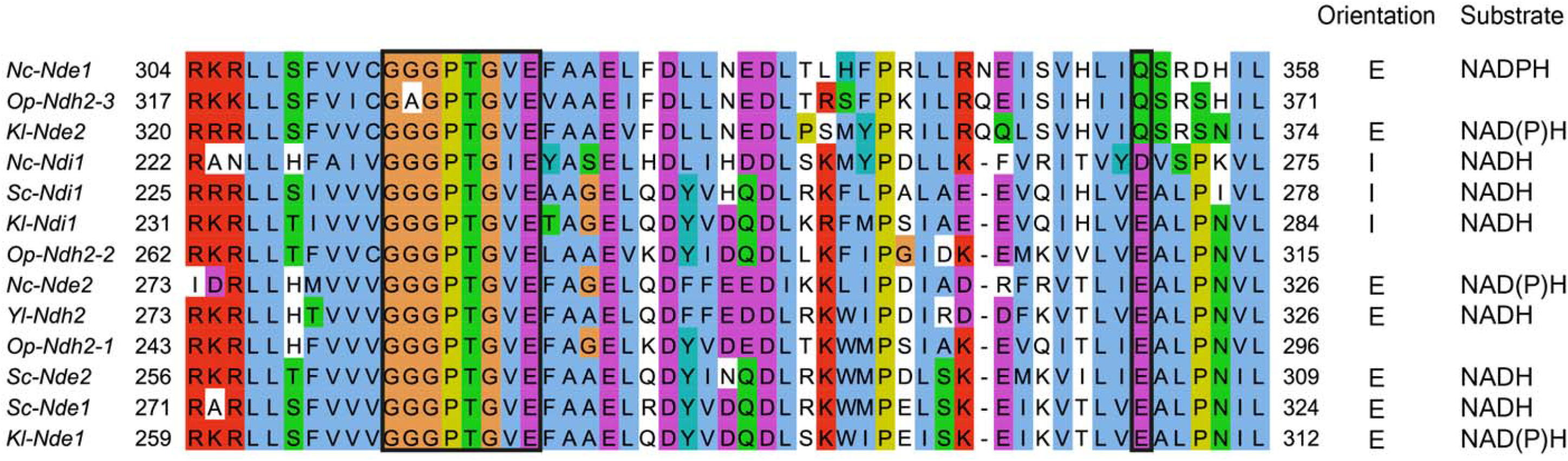
Section of MUSCLE sequence alignment of fungal/yeast NDH2 sequences, visualized by Jalview using Clustalx residue colors. Black boxes denote (left) conserved GxGxxGxE motif of second dinucleotide (substrate) binding domain and (right) residue affecting NADH/NADPH substrate specificity. Orientation, either I (internal) or E (external) and substrate utilization, NADH, NADPH or NAD(P)H (= both) are depicted for characterized fungal NDH2s. *Kl, Kluyveromyces lactis, Nc, Neurospora crassa, Op, Ogataea parapolymorpha, Sc, Saccharomyces cerevisiae, Yl, Yarrowia lipolytica*.

**Figure S2:**
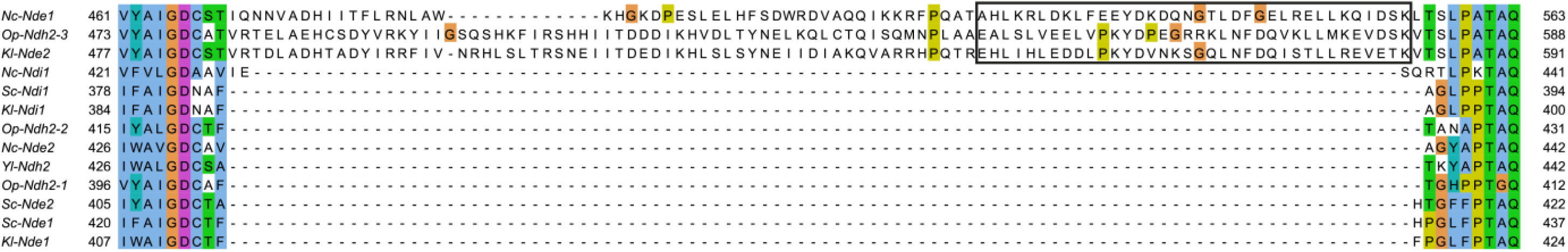
Section of MUSCLE multiple sequence alignment of fungal/yeast NDH2 sequences, visualized by Jalview using Clustalx residue colors. The black box denotes a (putative) EF-hand calcium binding domain as predicted by Motif Scan. *Kl, Kluyveromyces lactis, Nc, Neuurospora crassa, Op, Ogataea parapolymorpha, Sc, Saccharomyces cerevisiae*, Yl, *Yarrowia lipolytica*.

**Figure S3:**
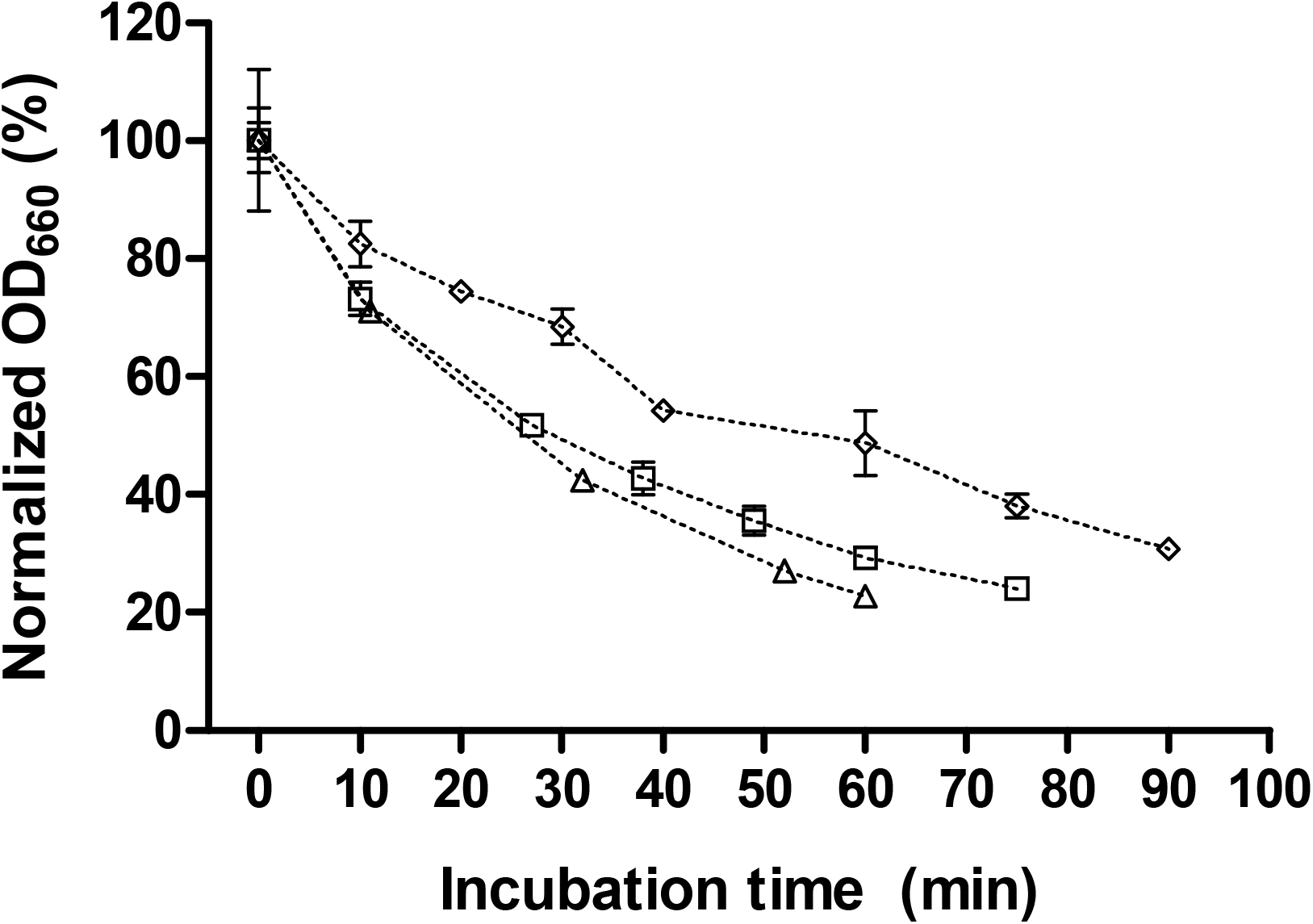
Proportion of osmotically insensitive cells of *O. parapolymorpha* strains CBS11895 (triangles), IMX2017 (diamonds) and IMX2197 (squares) during zymolyase treatment as determined by dilution in demineralized water. Data is presented as mean ± standard deviation from at least 2 independent replicates, and was normalized to initial OD for each strain individually.

**Table S1:** NAD(P)H oxidation (μmol (mg protein)^−1^ min^−1^) by *E. coli* membranes isolated from strains overexpressing individual *O. parapolymorpha* NDH2s. Control measurements were done with membranes isolated from a strain carrying an empty overexpression plasmid (pTrc99A). Assays were performed with a membrane protein concentration of 10 μg mL^−1^ at 37°C, with 200 μM (pH 7.4 and 8) or 100 μM (pH 5.5) NAD(P)H and 100 μM ubiquinone-1. For tests with calcium, 5 mM CaCl was used. Data is presented as mean ± standard deviation of at least duplicate measurements. ^a^Single measurement.

|  | NADH |  |  | NADPH |  |  |
| --- | --- | --- | --- | --- | --- | --- |
|  | pH 5.5 | pH 7.4 | pH 8.0 | pH 5.5 | pH 7.4 | pH 8.0 |
| Control<br>pTrc99A | $1.64 \pm 0.34$ | $1.64 \pm 0.34$ | $1.49 \pm 0.11$ | $0.03 \pm 0.04$ | $0.02 \pm 0.03$ | $0.18 \pm 0.01$ |
| Ndh2-1<br>pTrc99A-NDH2-1 | $4.47 \pm 0.05$ | $5.27 \pm 0.82$ | $4.58 \pm 0.25$ | < 0.01 | $0.03 \pm 0.05$ | $0.05 \pm 0.05$ |
| Ndh2-2<br>pTrc99A-NDH2-2 | $3.45 \pm 0.13$ | $3.95 \pm 0.85$ | $3.24 \pm 0.94$ | $0.06 \pm 0.08$ | $0.02 \pm 0.04$ | $0.09 \pm 0.09$ |
| Ndh2-3<br>pTrc99A-NDH2-3 | $3.77 \pm 0.05$ | $4.34 \pm 0.50$ | $4.70 \pm 0.02$ | $0.21 \pm 0.16$ | $0.01 \pm 0.02$ | $0.03 \pm 0.04$ |
| Control + $\text{Ca}^{2+}$<br>pTrc99A | 1.43<sup>a</sup> | $1.64 \pm 0.12$ | $1.64 \pm 0.09$ | $0.07 \pm 0.05$ | < 0.01 | $0.09 \pm 0.02$ |
| Ndh2-3 + $\text{Ca}^{2+}$<br>pTrc99A-NDH2-3 | $3.08 \pm 0.25$ | $4.15 \pm 0.69$ | $3.57 \pm 0.09$ | $0.17 \pm 0.16$ | < 0.01 | < 0.01 |

**Table S2:** Physiology of *Ogataea parapolymorpha* strain IMX2167 in aerobic, glucose-limited chemostat cultures grown at a dilution rate of 0.1 h^−1^ at 30°C and pH 5. Data are presented as mean ± mean absolute deviation from two independent replicates. Carbon recoveries were calculated based on a biomass carbon content of 48% (w/w). Symbols: Y_X/S_ and Y_X/O2_ = yield of biomass dry weight on glucose and oxygen, respectively; RQ = respiratory quotient; q_Glucose_, q_CO2_ and q_O2_ represent biomass-specific uptake/production rates of glucose, CO_2_ and O_2_, respectively; C_X_ represents biomass dry weight concentration.

| Strain | IMX2167 |
| --- | --- |
|  | *Δndh2-1 Δndh2-2 Δndh2-3 Δgut2* |
| Actual dilution rate (h<sup>-1</sup>) | 0.10 ± 0.00 |
| Reservoir glucose (g L<sup>-1</sup>) | 7.61 ± 0.01 |
| Y<sub>X/S</sub> (g biomass [g glucose]<sup>-1</sup>) | 0.52 ± 0.00 |
| Y<sub>X/O<sub>2</sub></sub> (g biomass [g O<sub>2</sub>]<sup>-1</sup>) | 1.44 ± 0.00 |
| RQ (-) | 1.05 ± 0.00 |
| q<sub>Glucose</sub> (mmol [g biomass]<sup>-1</sup> h<sup>-1</sup>) | -1.04 ± 0.00 |
| q<sub>CO<sub>2</sub></sub> (mmol [g biomass]<sup>-1</sup> h<sup>-1</sup>) | 2.25 ± 0.01 |
| q<sub>O<sub>2</sub></sub> (mmol [g biomass]<sup>-1</sup> h<sup>-1</sup>) | -2.14 ± 0.00 |
| C<sub>X</sub> (g biomass L<sup>-1</sup>) | 3.97 ± 0.00 |
| Carbon recovery (%) | 98.7 ± 0.3 |

**Table S3:** *S. cerevisiae* proteins (proposed to be) involved in ethanol-acetaldehyde, malate-oxaloacetate and malate-aspartate shuttle NADH shuttles, and corresponding ortholog proteins in *O. parapolymorpha*. Orthologs were identified using blastp (https://blast.ncbi.nlm.nih.gov/). *HPODL_03666 and HPODL_02528 have been experimentally verified as cytosolic and mitochondrial alcohol dehydrogenases in *O. parapolymorpha*, respectively (73, 74).

| *S. cerevisiae* protein | Annotation | *O. parapolymorpha* ortholog | E-value (blastp) |
| --- | --- | --- | --- |
| Adh1 | Cytosolic alcohol dehydrogenase | HPODL_03666* | 0.0 |
| Adh3 | Mitochondrial alcohol dehydrogenase | HPODL_02528* | 0.0 |
| Mdh1 | Mitochondrial malate dehydrogenase | HPODL_00710 | $2 \cdot 10^{-174}$ |
| Mdh2 | Cytosolic malate dehydrogenase | HPODL_02341 | $2 \cdot 10^{-83}$ |
| Aat1 | Mitochondrial aspartate aminotransferase | HPODL_01819 | $2 \cdot 10^{-91}$ |
| Aat2 | Cytosolic aspartate aminotransferase | HPODL_02844 | 0.0 |
| Dic1 | Mitochondrial dicarboxylate carrier | HPODL_04016 | $9 \cdot 10^{-112}$ |
| Oac1 | Mitochondrial oxaloacetate carrier | HPODL_00744 | $5 \cdot 10^{-157}$ |
| Odc1 / Odc2 | Mitochondrial oxodicarboxylate carrier | HPODL_01147 | $8 \cdot 10^{-144} / 3 \cdot 10^{-140}$ |
| Agc1 | Mitochondrial amino acid transporter | HPODL_03446 | $5 \cdot 10^{-122}$ |

**Table S4:**
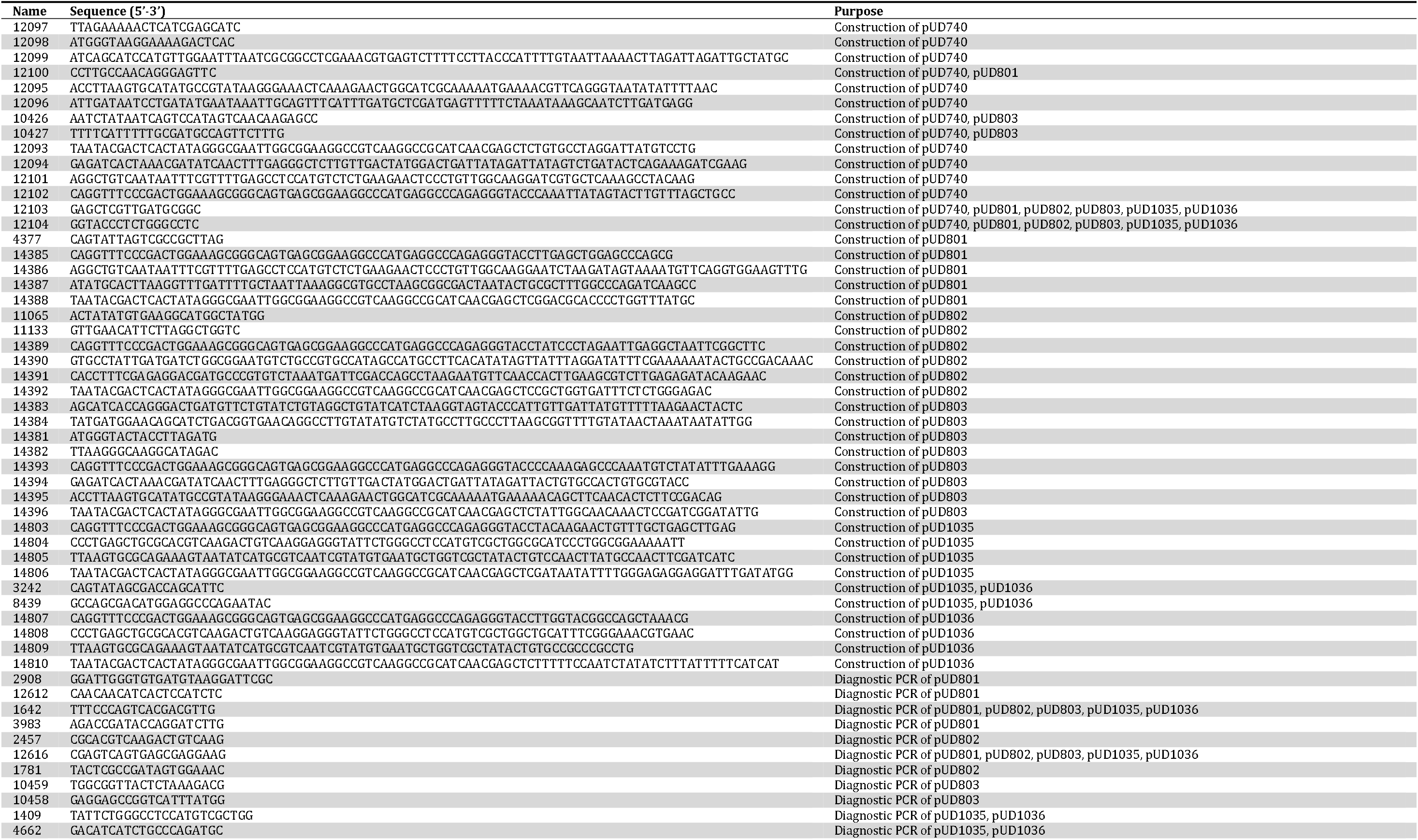
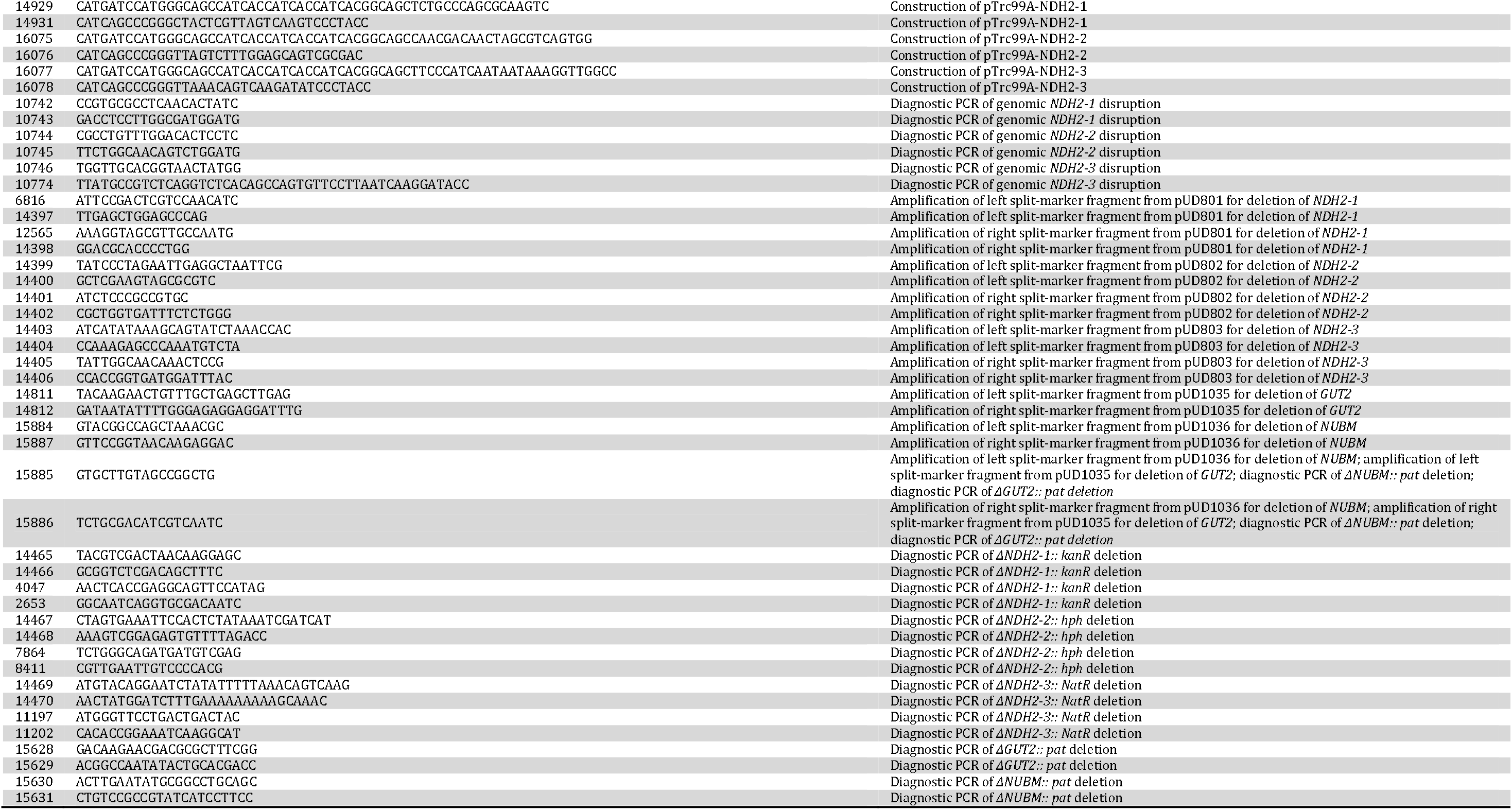
Primers used in this study.

